# A *C. elegans* genome-wide RNAi screen for altered levamisole sensitivity identifies genes required for muscle function

**DOI:** 10.1101/2020.12.01.407213

**Authors:** Timothy Chaya, Shrey Patel, Erin M. Smith, Andy Lam, Elaine N. Miller, Michael Clupper, Kirsten Kervin, Jessica Tanis

## Abstract

At the neuromuscular junction (NMJ), postsynaptic ionotropic acetylcholine receptors (AChRs) transduce a chemical signal released from a cholinergic motor neuron into an electrical signal to induce muscle contraction. To identify regulators of postsynaptic function, we conducted a genome-wide RNAi screen for genes required for proper response to levamisole, a pharmacological agonist of ionotropic L-AChRs at the *Caenorhabditis elegans* NMJ. A total of 117 gene knockdowns were found to cause levamisole hypersensitivity, while 18 resulted in levamisole resistance. Our screen identified conserved genes important for muscle function including some that are mutated in congenital myasthenic syndrome, congenital muscular dystrophy, congenital myopathy, myotonic dystrophy, and mitochondrial myopathy. Of the genes found in the screen, we further investigated those predicted to play a role in endocytosis of cell surface receptors. Loss of the Epsin homolog *epn-1* had opposing effects on the levels of postsynaptic L-AChRs and GABA_A_ receptors, resulting in increased and decreased abundance, respectively. This disrupts the balance of postsynaptic excitatory and inhibitory signaling, causing levamisole hypersensitivity. We also examined other genes that resulted in a levamisole hypersensitive phenotype when knocked down including *gas-1*, which functions in Complex I of the mitochondrial electron transport chain. Consistent with altered ATP synthesis impacting levamisole response, treatment of wild-type animals with levamisole resulted in L-AChR dependent depletion of ATP levels. These results suggest that the paralytic effects of levamisole ultimately lead to metabolic exhaustion.

## Introduction

Acetylcholine (ACh) released from motor neurons activates postsynaptic ionotropic ACh receptors (AChRs), resulting in an electrical signal that leads to muscle contraction. Disruption of cholinergic signaling at the neuromuscular junction (NMJ) is the underlying cause of severe muscle weakness observed in individuals with congenital myasthenic syndromes (CMS) and the autoimmune syndrome Myasthenia gravis (Engel *et al.* 2015). Clinical features of some congenital myopathies and muscular dystrophies resemble CMS despite different genetic causes and muscle histopathology (Mahjneh *et al.* 2013; Rodríguez Cruz *et al.* 2014; Montagnese *et al.* 2017). This suggests that characterizing mechanisms which regulate postsynaptic cholinergic signaling could provide insight into multiple neuromuscular disorders.

*C. elegans* body wall muscles are functionally comparable to vertebrate skeletal muscles and provide an excellent model for the study of neuromuscular transmission (Gieseler 2017). There are two classes of ionotropic AChRs at the *C. elegans* NMJ that have distinct pharmacological profiles. L-AChRs are heteromeric AChRs that are sensitive to the nematode-specific AChR agonist levamisole, while N-AChRs are homomeric AChRs that are activated by nicotine, but insensitive to levamisole (Richmond and Jorgensen 1999; Touroutine *et al.* 2005). Wild-type animals exposed to levamisole undergo time-dependent paralysis (Lewis *et al.* 1980). Altered sensitivity to levamisole-induced paralysis can be indicative of defects in postsynaptic cholinergic signaling. A forward genetic screen for mutants with strong levamisole resistance identified mutations in 12 different genes encoding subunits of the L-AChR as well as proteins required for L-AChR trafficking, clustering, and muscle contraction (Lewis *et al.* 1980; Gally *et al.* 2004; Eimer *et al.* 2007; Gendrel *et al.* 2009). Levamisole resistant mutants have also been isolated in a forward genetic screen for animals that initially exhibit paralysis in response to levamisole, but subsequently adapt and regain motility (Rapti *et al.* 2011; Boulin *et al.* 2012; Richard *et al.* 2013; D’alessandro *et al.* 2018). However, mutants hypersensitive to levamisole, as well as levamisole resistant mutants which also have a lethal or sterile phenotype, could not be isolated in these genetic screens. For example, loss of function mutations in *fer-1*, the *C. elegans* homolog of Dysferlin, which is mutated in Limb-Girdle Muscular Dystrophy 2B, result in weak levamisole resistance, but also sterility (Achanzar and Ward 1997; Krajacic *et al.* 2013). Since *fer-1* was not identified in the forward genetic screens for levamisole resistance, this suggests that performing a different type of genetic screen may uncover new genes important for muscle function.

Feeding *C. elegans* bacteria that make double-stranded RNA (dsRNA) produces an RNA interference (RNAi) signal that spreads throughout the animal, causing knockdown of corresponding gene function (Timmons *et al.* 2001). Since animals treated with dsRNA do not have to be propagated and the dsRNA can be delivered post-embryonically, genome-wide RNAi screens can identify genes that are otherwise essential for viability. Prior RNAi screens that assessed approximately 10% of the genome for resistance and hypersensitivity to the acetylcholine esterase inhibitor aldicarb identified genes important for cholinergic neurotransmission and GABAergic signaling, respectively (Sieburth *et al.* 2005; Vashlishan *et al.* 2008). Here we performed a systematic genome-wide RNAi screen for gene knockdowns that cause hypersensitivity or resistance to levamisole to identify novel regulators of postsynaptic signaling. We identified 135 gene knockdowns that resulted in altered levamisole response. Only 7% of the genes that we discovered had a previously annotated levamisole phenotype, suggesting that our screen identified new regulators of postsynaptic function.

## Materials and Methods

### Nematode culture

*C. elegans* were maintained on Nematode Growth Medium (NGM) plates with OP50 *E. coli* using standard techniques (Brenner 1974). The wild-type strain was Bristol N2. Other strains used in this study were as follows: CB1072 *unc-29(e1072)* I, ZZ17 *lev-10(x17)* I, CB407 *unc-49(e407)* III, GR1373 *eri-1(mg366)* IV, CW152 *gas-1(fc21)* X, GS2526 *arIS37* I; *mca-3(ar492) dpy-20(e1282)* IV, UDE5 *unc-63(kr98-YFP)* I; *eri-1(mg366)* IV, UDE23 *krSi2(unc-49::tagRFP)* III; *eri-1(mg366)* IV. All *C. elegans* strains were cultured at 20°C except those with *eri-1(mg366)* which were maintained at 15°C.

### Levamisole time course assay

24 well NGM plates or RNAi plates (NGM plus 1 mM IPTG and 25 μg/μl carbenicillin) with 40 μl of bacterial food source per well were prepared. In each plate, there were at least three independent wells per genotype (Figures 1A, 4F) or RNAi clone (Figures 2A,B and 4A-E); researchers who performed the assay were blinded. Approximately twenty first larval stage (L1) animals synchronized by bleaching were pipetted into each well. After three days, 0.4 mM levamisole (Sigma) in M9 buffer was added to wells at intervals; the number of moving worms per well was counted every 5 minutes until all worms stopped moving or 60 minutes passed. The total number of worms per well was recorded at the end of the assay. Data were used to generate Kaplan-Meier survival plots in Graph Pad Prism and Log-rank (Mantel-Cox) tests were used to determine significance.

**Figure 1:**
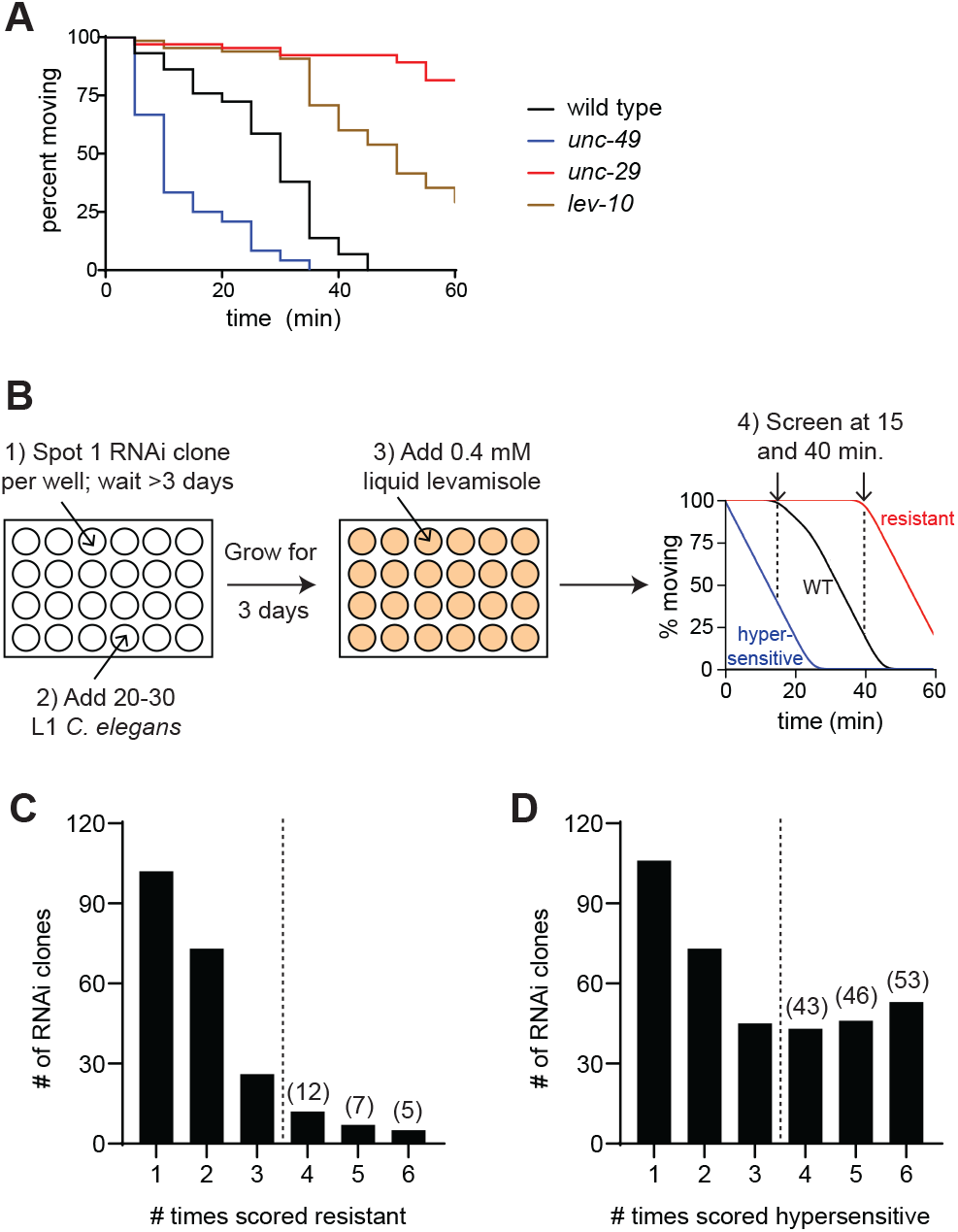
RNAi screen for altered sensitivity to levamisole. (A) Time-dependent paralysis of wild-type (black), *unc-29* (red), *lev-10* (tan) and *unc-49* (blue) animals in a levamisole swim assay. Loss of *unc-29* (red) and *lev-10* (tan) caused resistance; loss of *unc-49* (blue) resulted in hypersensitivity. n≥30 per genotype in this representative experiment; p <0.0001 for all mutants compared to the wild-type control. (B) Workflow for the RNAi screen. *eri-1* L1s were fed bacteria expressing dsRNA in 24 well plates. After three days, 0.4 mM levamisole in M9 was added to each well and animals were scored for hypersensitivity after 15 minutes and resistance after 40 minutes. (C-D) After the primary screen was performed in duplicate, “hits” were screened again in quadruplicate. Gene knockdowns were considered positive if altered levamisole sensitivity was observed four or more times (p < 0.05).

**Figure 2:**
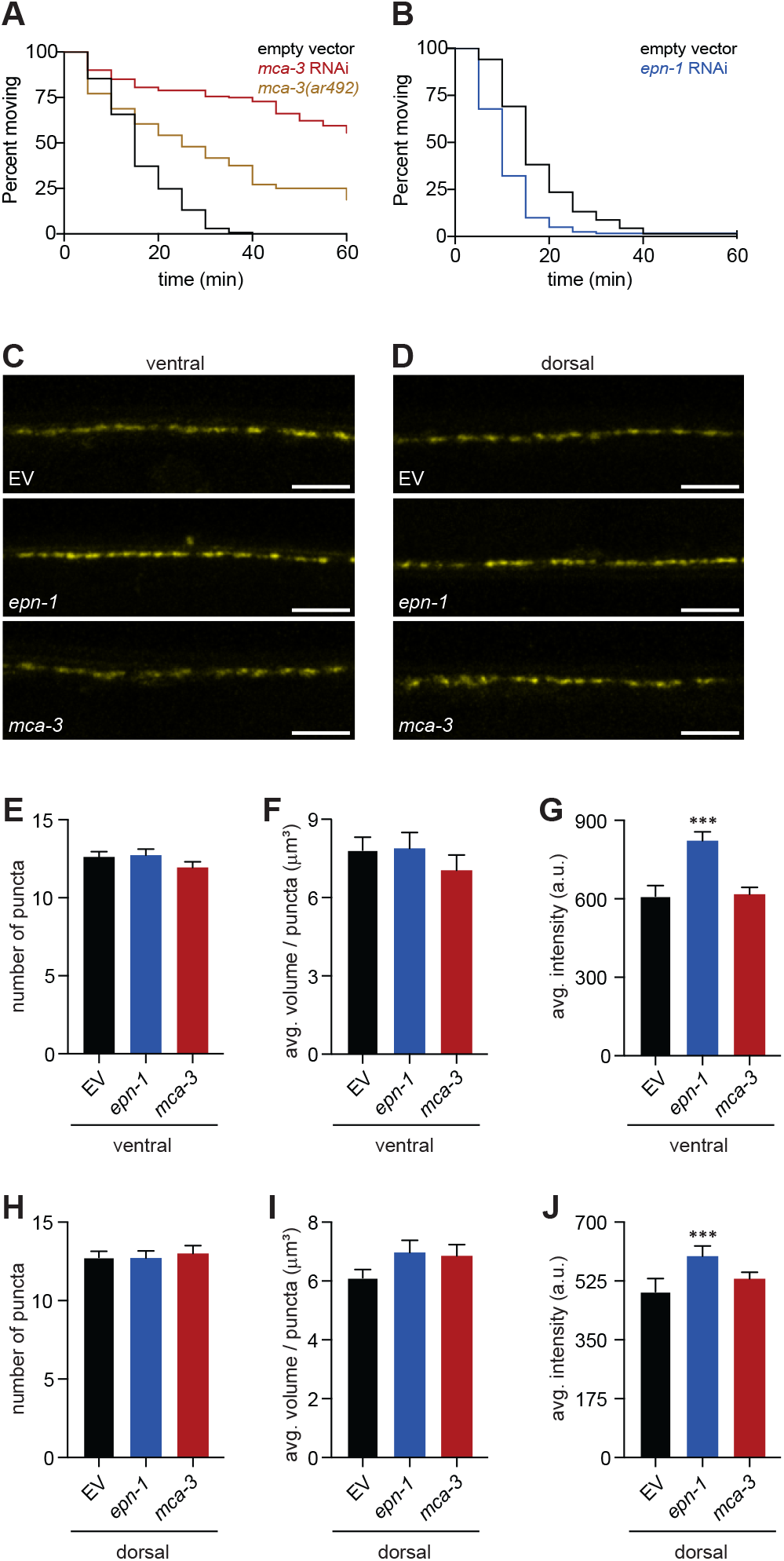
Loss of EPN-1 increases the abundance of L-AChRs. (A) *mca-3* knockdown (red) and the *mca-3(ar492)* mutation (tan) both cause resistance to levamisole induced paralysis; p< 0.0001, n ≥ 50. (B) Knockdown of *epn-1* (blue) results in hypersensitivity to levamisole; p < 0.0001, n ≥ 68 (C,D) Representative images of YFP-tagged UNC-63 L-AChRs on the ventral (C) and dorsal (D) sides of animals exposed to EV (top), *epn-1* (middle), and *mca-3* (bottom) RNAi. (E-J) Quantitation of the density (E,H), volume (F,I), and abundance (G,J) of UNC-63::YFP clusters at the NMJ in EV, *epn-1*, and *mca-3* knockdown animals; ventral (E-G) and dorsal (H-J) measurements are indicated. Data are mean ± SE, n ≥ 16, *** p < 0.001.

### Levamisole sensitivity RNAi screen

The RNAi feeding screen was carried out in 24 well format as described (Kamath *et al.* 2003). Approximately thirty L1 *eri-1(mg366)* animals were deposited in each well. After three days of RNAi exposure at 20°C, wells were checked for the presence of adult worms and lack of contamination. 0.4 mM levamisole (Sigma) in M9 buffer was added and the number of moving animals in each well was recorded at 15 and 40 minutes to identify hypersensitivity and resistance, respectively. Wells with at least 50% of the animals paralyzed at 15 minutes were marked as hypersensitive, while wells with at least 25% of the animals moving at 40 minutes were scored as resistant. Animals grown on the HT115 empty vector (EV) control were scored every day of screening. The entire genome-wide screen was performed in duplicate. Knockdowns that caused altered levamisole response twice, or once for genes with known muscle expression, were retested in quadruplicate. For the final gene list (Supplemental Table 1), the RNAi clones caused a levamisole phenotype in at least 4/6 trials, did not inhibit mobility in a 60-minute swim test in water, produced a levamisole phenotype in a time course assay, and were sequence verified.

### Fluorescence microscopy and image analysis

UDE5 *unc-63(kr98-YFP)*; *eri-1(mg366)* animals to visualize L-AChRs, and UDE23 *krSi2(unc-49::tagRFP)*; *eri-1(mg366)* animals to visualize GABA_A_ receptors (GABA_A_Rs), were grown on EV, *epn-1* and *mca-3* RNAi plates. Late fourth larval stage (L4) stage animals were picked onto new corresponding RNAi plates, cultured for 24 hours at 20° to produce staged adults, and immobilized with 50 mM sodium azide on 3% agar pads ten minutes prior to imaging. UNC-63::YFP Z-stack images were obtained with a Zeiss LSM880 confocal microscope using a 40X water objective and master gain of 870. UNC-49::tagRFP images were collected with a Zeiss LSM780 confocal microscope using a 40X water objective and master gain of 981. Z-stack images were deconvoluted using Huygens Professional Software with signal to noise ratio set to 5 and quality threshold of 0.05. Images with no indication of movement were analyzed with Volocity software with a minimum object size of 1 μm^3^. The number of puncta per 50 μm segment, volume (μm^3^) of each punctum, and mean intensity (arbitrary units) of each punctum were recorded. Welch’s two tailed t-test with unequal variance was used to test for significance.

### ATP quantitation

ATP quantification assays were performed with the ATP Bioluminescence Assay Kit CLS II (Roche Diagnostics) as described (Palikaras and Tavernarakis 2016) with some modifications. Synchronized populations of *unc-29(e1072)* and wild-type animals, ~1000 per 10 cm NGM plate, were exposed to the following conditions: 60 minutes no treatment, 60 minutes M9, or 60 minutes 0.4 mM levamisole in M9. Animals were washed 3x with M9 solution and 20 μl of worms were transferred to microcentrifuge tubes. 180 μl of boiling Tris-EDTA buffer (100mM Tris, 4mM EDTA pH 7.75) was added and samples were incubated at 100°C for 2 minutes, sonicated for 4 minutes (Model 150V/T Ultrasonic Homogenizer), and then centrifuged at 14,000 RPM for 10 minutes at 4°C. 30 μl of the supernatant was added to 270 μl Tris-EDTA buffer to make 1:10 dilutions for triplicate technical replicates. ATP content was determined following the Roche ATP Bioluminescence Assay Kit CLS II protocol using a Glomax 96 Microplate Luminometer. ATP levels were normalized to total protein content (Pierce BCA protein assay kit). Six biological replicates were performed; statistical significance was determined using a non-parametric one-way ANOVA.

#### Data availability

All data and methods required to confirm the conclusions of this work are within the article, figures, and table. Table S1 is a list of all genes identified in the RNAi screen that were confirmed by time course assays and sequencing. This supplemental material is available at figshare.

## Results and Discussion

### An RNAi screen for altered sensitivity to levamisole

The body wall muscles receive excitatory and inhibitory inputs from cholinergic and GABAergic motor neurons, respectively. Altered time to levamisole-induced paralysis can be used to identify genes that impact the balance of postsynaptic excitatory and inhibitory signaling. While levamisole resistant mutants have been isolated, mutations in genes required for both viability and levamisole response could not be identified in the forward genetic screens, and there has never been a genetic screen for levamisole hypersensitivity (Lewis *et al.* 1980; Rapti *et al.* 2011; Boulin *et al.* 2012; Richard *et al.* 2013; D’alessandro *et al.* 2018). We reasoned that a genome-wide RNAi screen for gene knockdowns that cause hypersensitivity or resistance to levamisole would identify new regulators of postsynaptic function.

Traditionally, resistance or hypersensitivity to levamisole has been determined using a plate-based assay which requires prodding animals to assess paralysis. To enable high-throughput screening, we developed a liquid levamisole swim assay performed on animals grown in 24 well plates. Compared to plate-based experiments, the liquid levamisole assay required a lower concentration of levamisole, 0.4 mM, to induce complete paralysis of wild-type animals within 45 minutes (Figure 1A). Further, the rigorous swimming of the animals in liquid allowed us to assess paralysis without physically prodding the animals. We tested mutants with known levamisole phenotypes in our new assay and found that mutations in *unc-29*, which encodes an essential L-AChR subunit and *lev-10*, which is important for L-AChR clustering, caused resistance (Figure 1A) (Lewis *et al.* 1980; Gally *et al.* 2004). The swim assay also detected hypersensitivity to levamisole in the *unc-49* GABA_A_R mutant, which has reduced inhibitory signaling (Figure 1A) (Richmond and Jorgensen 1999; Bamber *et al.* 2005). These results demonstrate that levamisole phenotypes can be observed using a swim-based levamisole assay and since this assay is less labor intensive, it is ideal for a genetic screen.

We performed liquid levamisole swim assays on animals grown on RNAi clones in 24-well plates and scored the number of animals moving at two time points to identify hypersensitivity and resistance (Figure 1B). In the primary screen, we tested animals fed clones from the *C. elegans* RNAi library generated by the Ahringer group (Kamath *et al.* 2003), representing ~86% of the genome, in duplicate. Gene knockdowns that caused altered levamisole response twice, or once for genes with known muscle expression, were defined as ‘hits’ and retested in quadruplicate. We identified 142 RNAi clones that caused hypersensitivity and 24 RNAi clones that produced resistance in at least four out of six trials and did not inhibit mobility during a 60-minute swim test in water, which ensured that these animals were capable of movement for the entire assay time period (Figure 1C, D). 93% of the gene knockdowns that caused levamisole hypersensitivity in the screen paralyzed significantly faster than the control in time course experiments; 75% of the gene knockdowns that were found to elicit resistance in the screen were validated with time course assays. In total, we sequence-verified 135 of the RNAi clones that caused altered levamisole sensitivity in time course assays (Supplemental Table 1). 89% have human homologs, and of those with a defined expression pattern, 74% are muscle expressed. This indicates that we identified highly conserved genes with postsynaptic expression at the NMJ.

Just 7% of the genes that we discovered were previously associated with altered levamisole sensitivity. Of the RNAi clones for nine of the genes identified in the forward genetic screen for strong levamisole resistance, we identified only a third of these, *unc-63*, *lev-10* and *unc-22*, in our screen. While many other genes that impact levamisole sensitivity were also not identified in our screen, there are multiple possible reasons for this. First, not all of the RNAi clones in the library cause efficient knockdown. Second, since RNAi exposure commenced at the first larval stage, there may have been insufficient knock down of some genes to generate an altered levamisole response by the time the animals were assayed. Third, some of the RNAi clones caused severe developmental defects that prevented us from screening the levamisole sensitivity of those animals. Fourth, we used a different levamisole assay than the forward genetic screens and thus we were unable to identify animals that initially exhibit paralysis in response to levamisole, but subsequently adapt and regain motility (Rapti *et al.* 2011; Boulin *et al.* 2012; Richard *et al.* 2013; D’alessandro *et al.* 2018). Finally, contamination and lack of animals in a limited number of wells prevented us from screening some library clones in duplicate. Despite these limitations, our screen identified 126 new genes that exhibit a levamisole phenotype when knocked down.

Our reverse genetic screen provided unique advantages that enabled us to discover these new genes required for proper levamisole response. We were able to detect both hypersensitive and resistant phenotypes, performed an unbiased screen using the entire *C. elegans* RNAi feeding library rather than cherry picking clones, and initiated gene knockdown after embryogenesis was complete, which allowed us to identify many genes otherwise essential for viability. Notably, 50% of the genes that we discovered have an annotated sterile or lethal phenotype, justifying the need for a reverse genetic screen to identify additional factors that impact levamisole response. Here, we have performed an initial characterization of two different classes of genes identified in our screen.

### *C. elegans* Epsin regulates postsynaptic receptor abundance

During endocytosis, the plasma membrane and associated proteins are pulled toward the cytosol, membrane scission occurs, and the endocytic vesicle is internalized. Cargoes are then either targeted for degradation or recruited back to the plasma membrane via recycling endosomes to modulate synaptic strength (Park *et al.* 2004; Wang *et al.* 2008). Maintenance of the proper abundance of postsynaptic receptors is achieved by balancing the delivery of new receptors to the plasma membrane with the endocytic removal of existing receptors. Genes required for L-AChR assembly, trafficking to the cell surface, and clustering on the body wall muscles, but not internalization have been previously identified (Gally *et al.* 2004; Eimer *et al.* 2007; Gendrel *et al.* 2009; Rapti *et al.* 2011; Richard *et al.* 2013; D’alessandro *et al.* 2018). We discovered two genes in our screen, *mca-3* and *epn-1*, which have previously described roles in endocytosis. MCA-3, a plasma membrane Ca^2+^ ATPase (PMCA), functions in clathrin-mediated endocytosis in *C. elegans* coelomocytes by recruiting endocytic machinery to the plasma membrane (Bednarek *et al.* 2007). Further, in hippocampal neurons, inhibition of PMCA2 triggers the loss of ionotropic α7-nAChRs (Gómez-Varela *et al.* 2012). Knockdown of *mca-3* caused levamisole resistance in time course assays (Figure 2A). Although null mutations in *mca-3* cause lethality, we were able to validate our results with a partial loss of function mutation and found that this generated significant levamisole resistance, though a weaker phenotype than observed with *mca-3* knockdown (Figure 2A). EPN-1 is a homolog of mammalian Epsin, which functions as an adaptor to recruit specific cargoes and induce membrane curvature that results in endocytic vesicle formation (Chen *et al.* 1998; Wendland *et al.* 1999; Ford *et al.* 2002; Wang *et al.* 2006). Knockdown of *epn-1* resulted in levamisole hypersensitivity, though we could not confirm this result with an *epn-1* mutant as all known mutants are non-viable (Figure 2B). Both *mca-3* and *epn-1* are expressed in the body wall muscles where they could play a role in regulating the abundance of postsynaptic receptors (Bednarek *et al.* 2007; Shen *et al.* 2013).

To determine if loss of *mca-3* or *epn-1* had an impact on L-AChR abundance we used a knock-in strain that expresses the YFP-tagged UNC-63, an essential subunit of the L-AChR, from its genomic locus (Gendrel *et al.* 2009) and analyzed the distribution of UNC-63::YFP puncta in animals grown on EV, *mca-3*, and *epn-1* RNAi clones. We quantitated the number of puncta per 50 μm to define the quantity of postsynaptic specializations, the average volume of each puncta to determine the size of these specializations, and the average intensity to measure the relative number of receptors. Based on these parameters, loss of *mca-3* did not affect L-AChR abundance or localization (Figure 2C-J). Therefore, the levamisole resistant phenotype associated with loss of *mca-3* cannot be explained by a reduced number of L-AChRs. However, loss of *epn-1* caused a significant increase in UNC-63::YFP intensity, without altering puncta number or size, on both the dorsal and ventral sides of the animals (Figure 2C-J). This increase in the number of cell surface L-AChRs is consistent with the levamisole hypersensitive phenotype observed in *epn-1* knockdown animals. Our results suggest that EPN-1 plays a role in the endocytosis of L-AChRs from the plasma membrane without affecting the number or size of postsynaptic specializations.

Defects in GABA signaling can disrupt the excitatory-inhibitory balance in the body wall muscles, resulting in altered levamisole response (Vashlishan *et al.* 2008; Krajacic *et al.* 2013; Kowalski *et al.* 2014). We reasoned that an increase in postsynaptic UNC-49 GABA_A_Rs and thus inhibitory signaling, could lead to a levamisole resistant phenotype such as that observed with loss of *mca-3*. In fact, *cup-5*, a mutant identified along with *mca-3* (*cup-7*) in a forward genetic screen for coelomocyte uptake defective mutants regulates plasma membrane levels of GABA_A_Rs (Fares and Greenwald 2001; Davis *et al.* 2010). Using a transgenic strain that expresses RFP-tagged UNC-49 at single copy (Pinan-Lucarré *et al.* 2014), we analyzed the distribution of postsynaptic UNC-49::tagRFP receptor puncta in animals grown on EV, *mca-3*, and *epn-1* RNAi clones. We found that, for the most part, loss of *mca-3* did not alter GABA_A_R abundance or localization, though a small, yet significant decrease in the average volume of GABA_A_R puncta was observed on the dorsal side (Figure 3A-H). Surprisingly, analysis of GABA_A_Rs in the *epn-1* knockdown showed a significant decrease in UNC-49::tagRFP intensity compared to the control, with no effect on the number or size of the puncta (Figure 3A-H). These results suggest that EPN-1 regulates the abundance of both L-AChRs and GABA_A_Rs at the neuromuscular junction to maintain the appropriate balance of excitatory and inhibitory signaling.

**Figure 3:**
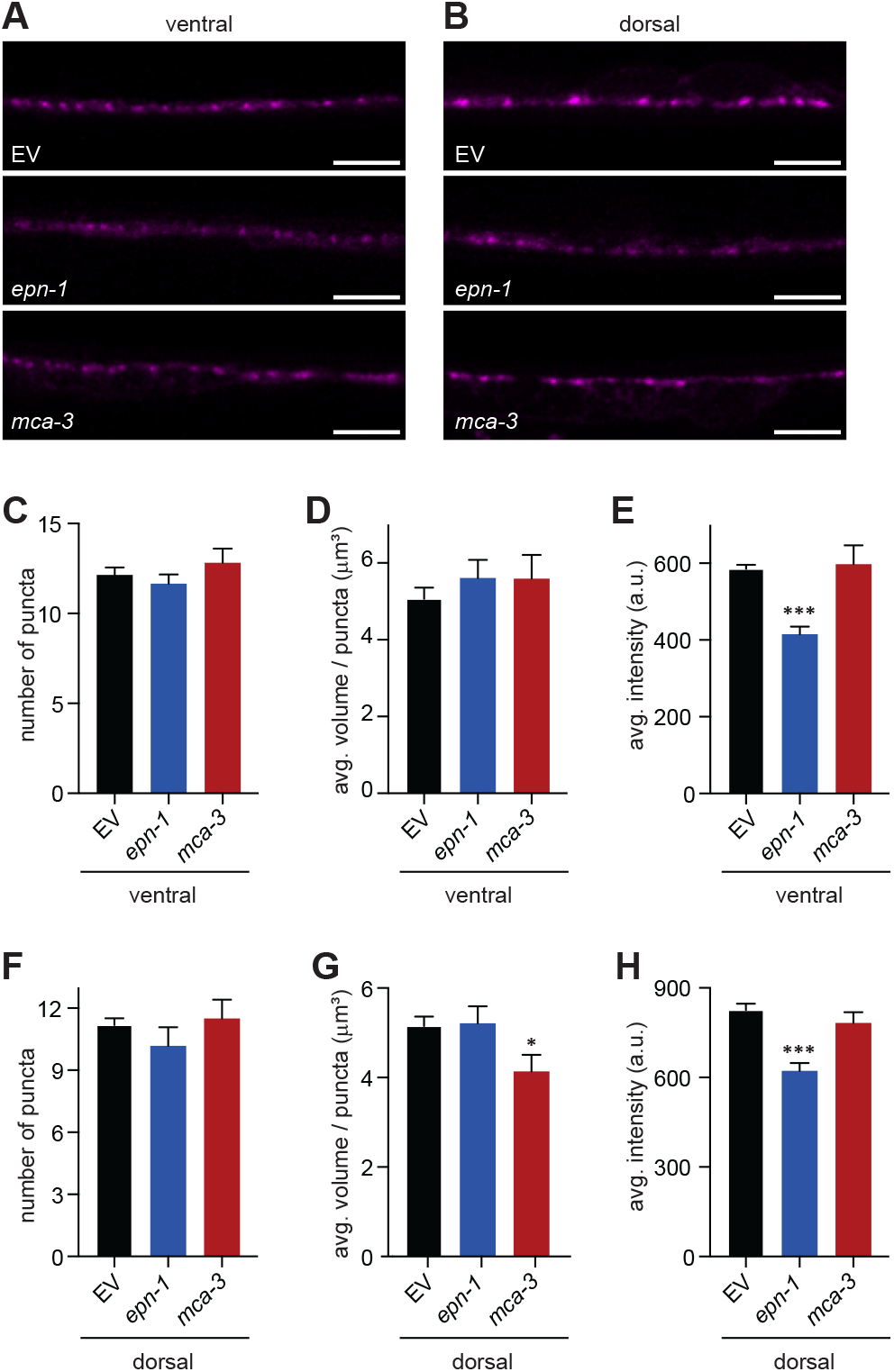
Loss of EPN-1 decreases postsynaptic GABA_A_R abundance. (A,B) Representative images of RFP-tagged UNC-49 GABA_A_Rs on the ventral (A) and dorsal (B) sides of animals exposed to EV (top), *epn-1* (middle), and *mca-3* (bottom) RNAi. (C-H) Quantitation of UNC-49::tagRFP density (C,F), volume (D,G), and abundance (E,H) on the body wall muscles of EV, *epn-1*, and *mca-3* knockdown animals; ventral (C-E) and dorsal (F-H) measurements are indicated. Data are mean ± SE, n ≥ 17, * p < 0.05, *** p < 0.001.

Both the increase in L-AChRs and decrease in GABA_A_Rs on the body wall muscles likely contribute to the levamisole hypersensitive phenotype observed in *epn-1* knockdown animals. However, the mechanism by which EPN-1 has opposing effects on the abundance of two different types of postsynaptic receptors, impacting both excitatory and inhibitory synapses, is not known. EPN-1 has been shown to be expressed in multiple cell types including body wall muscles and neurons (Shen *et al.* 2013). As loss of *epn-1* increases L-AChR abundance, EPN-1 may act directly in the body wall muscles to regulate the endocytosis of these receptors. How EPN-1 controls the levels of GABA_A_Rs is likely less straightforward. Mutants with GABA transmission defects can display levamisole hypersensitivity (Vashlishan *et al.* 2008; Kowalski *et al.* 2014). Further, the *C. elegans* Punctin MADD-4B, secreted by GABAergic neurons, is required for appropriate recruitment of GABA_A_Rs to the NMJ (Pinan-Lucarré *et al.* 2014; Tu *et al.* 2015). Thus, it is possible that EPN-1 plays an endocytic role in GABAergic neurons, leading to a compensatory effect on postsynaptic GABA_A_Rs. Future studies will define the EPN-1 site of action, subcellular distribution, and effect on other synaptic markers as well as whether postsynaptic receptor abundance is regulated by clathrin-mediated or clathrin-independent endocytosis.

### Disruption of cellular ATP levels alters levamisole sensitivity

Although levamisole has been used for decades to identify mutants with defects in postsynaptic signaling and immobilize animals for microscopy, the mechanism by which treatment with this pharmacological agent results in paralysis and death has not been fully defined. L-AChR activation leads to opening of the voltage-gated Ca^2+^ channel EGL-19, muscle depolarization, and ultimately, Ca^2+^ release from intracellular stores through the ryanodine receptor UNC-68 (Liu *et al.* 2011). Ca^2+^ binds to troponin, causing a shift in the position of tropomyosin and increasing the probability that myosin will bind the thin filament to produce force to generate muscle contraction. The sarco-endoplasmic reticulum Ca^2+^ ATPase (SERCA) works in concert with plasma membrane Ca^2+^ ATPases (PMCAs) and Na^+^/Ca^2+^ exchangers to lower cytoplasmic Ca^2+^ levels, which terminates contraction (Clapham 2007; Martin and Richmond 2018). If there is a decline in cellular ATP levels, this prevents Ca^2+^ from being pumped out of the cytoplasm, causing the muscles to remain contracted until Ca^2+^ dependent proteases induce muscle degeneration and finally relaxation in death (Galimov *et al.* 2018). It has been hypothesized levamisole-induced paralysis and death could be the result of cellular metabolic catastrophe, a condition in which ATP depletion prevents muscle function (Lewis *et al.* 1980). We discovered that knockdown of 14 genes predicted to play a role in ATP synthesis, *gas-1*, *cox-7c*, c*chl-1*, *nduf-9*, *cox-4*, *nuo-3*, C33A12.1, *cyc-2.1*, *yars-2*, *hpo-18*, *asg-1*, F53F4.10, F42G8.10 and D2030.4, caused levamisole hypersensitivity both in our genetic screen and time course assays (Figure 4A-E). While most of these genes are essential for viability (Supplemental Table 1), we were able to validate the levamisole hypersensitivity phenotype with a *gas-1* loss of function mutant, which has defects in Complex I of the mitochondrial electron transport chain (Figure 4F) (Kayser *et al.* 2001). These results suggest that reduced ATP synthesis increases sensitivity to levamisole, accelerating death.

**Figure 4:**
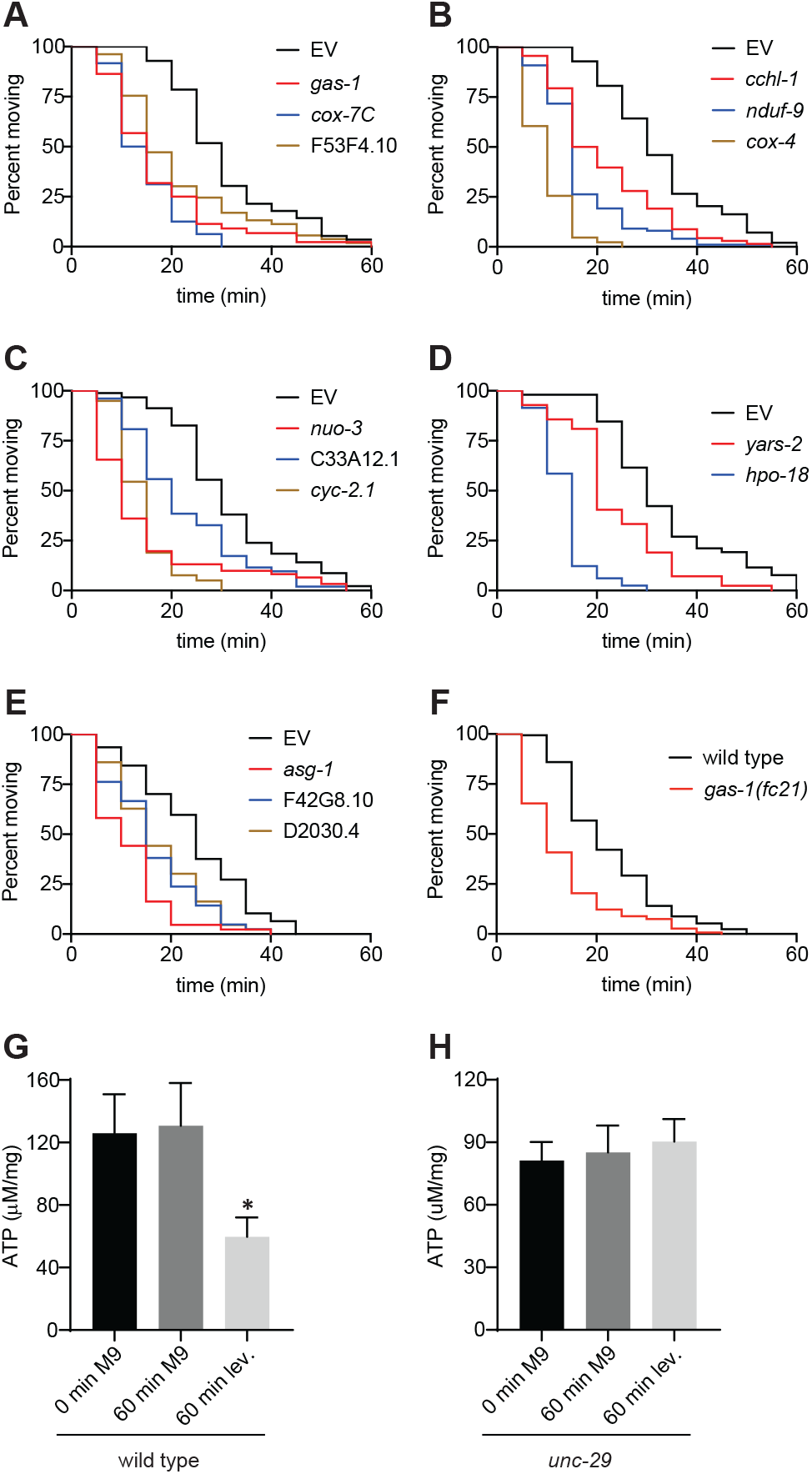
Levamisole exposure reduces cellular ATP. (A-E) A functional class with genes required for ATP synthesis was identified in the RNAi screen for altered sensitivity to levamisole. Knockdown of *gas-1* (red), *cox-7c* (blue) and F53F4.10 (tan) in (A), c*chl-1* (red), *nduf-9* (blue) and *cox-4* (tan) in (B), *nuo-3* (red), C33A12.1 (blue), and *cyc-2.1* (tan) in (C), *yars-2* (red) and *hpo-18* (blue) in (D), and *asg-1* (red), F42G8.10 (blue) and D2030.4 (tan) in (E) causes hypersensitivity to levamisole compared to EV (black) in time course assays; p < 0.001 for all knockdowns. (F) Like the *gas-1* knockdown in (A), *gas-1(fc21)* (red) exhibits hypersensitivity to levamisole induced paralysis compared to the wild type (black); p < 0.001. (G,H) 60 minutes of exposure to 0.4 mM levamisole (lev) causes a significant decrease in ATP (* p < 0.05) in wild type (G), but not *unc-29* L-AChR mutants (H); data are the mean ± SE of six independent experiments.

To determine if levamisole exposure leads to metabolic exhaustion, we quantitated ATP levels in animals that were washed off plates, swimming in M9 buffer for 60 minutes, or swimming in 0.4 mM levamisole for 60 minutes. While swimming in buffer had no effect on measured ATP, addition of levamisole significantly decreased ATP levels (Figure 4G). This was due to signaling through L-AChRs since levamisole did not have an impact on ATP levels in the *unc-29* L-AChR mutant (Figure 4H). This shows that levamisole exposure decreases ATP in cells that express L-AChRs and provides evidence for levamisole-induced metabolic catastrophe.

It is possible that knockdown of some seemingly unrelated genes identified in our screen may result in hypersensitivity to levamisole due to reduced cellular ATP and future studies will test this hypothesis. Loss of *pat-4* (Integrin Linked Kinase), *tln-1* (Talin I), *unc-73* (Trio), and *unc-60* (Cofilin) causes fragmentation of the muscle mitochondrial network, while loss of *prmt-1* (Protein Arginine Methyltransferase I) has been shown to reduce ATP synthesis (Etheridge *et al.* 2015; Sha *et al.* 2017). Likewise, we reason that mutants with reduced ATP usage would produce a resistant phenotype. SERCA and PMCA together utilize >30% of muscle produced ATP to pump Ca^2+^ out of the cytoplasm (Homsher 1987). Since release of Ca^2+^ from intracellular stores through ryanodine receptors dictates the amount of ATP used by Ca^2+^ ATPases, ATP usage can be attenuated by decreasing Ca^2+^ release from intracellular stores. This provides a possible explanation for the levamisole resistant phenotype observed in the *unc-68* ryanodine receptor mutant (Lewis *et al.* 1980). SERCAs translocate two Ca^2+^ ions into the sarco-endoplasmic reticulum lumen for each molecule of ATP hydrolyzed, while PMCAs translocate a single Ca^2+^ ion per ATP (Inesi and De Meis 1989; Hao *et al.* 1994). Thus, loss of the PMCA MCA-3 may reduce ATP usage, enabling the more efficient SERCA SCA-1 to continue to pump Ca^2+^ into the sarco-endoplasmic reticulum and leading to the levamisole resistance observed in the *mca-3* mutant. In conclusion, our results suggest that levamisole activation of L-AChRs causes excessive release of Ca^2+^ from intracellular stores, which disrupts cellular utilization of ATP levels. Consequently, loss of genes that affect either ATP synthesis or usage impacts levamisole sensitivity.

## Conclusion

Here we conducted the first genome-wide reverse genetic screen for postsynaptic regulators. In summary, we identified 135 gene knockdowns that caused altered sensitivity to the L-AChR agonist levamisole, describing a levamisole phenotype for 93% of these genes for the first time. We discovered novel genes because this was the first genetic screen capable of detecting levamisole hypersensitivity and genes that impact both levamisole response and viability. One gene that causes levamisole hypersensitivity when knocked down and lethality when knocked out is *epn-1*. Our work shows that loss of *epn-1* impacts postsynaptic function by having opposing effects on the abundance of postsynaptic L-AChRs and GABA_A_Rs. Notably, our screen also identified *C. elegans* homologs of genes mutated in congenital myasthenic syndrome (AChR), congenital muscular dystrophy (Mannose-1-phosphate guanyltransferase beta), congenital myopathy (Cofilin), myotonic dystrophy (ELAV-type RNA binding protein), and mitochondrial myopathy (NADH dehydrogenase) (Lu *et al.* 1999; Milne and Hodgkin 1999; Agrawal *et al.* 2007; Ockeloen *et al.* 2012; Carss *et al.* 2013; Engel *et al.* 2015; Schubert and Vilarinho 2020). This suggests that altered response to levamisole can be used broadly to discover and study genes that impact muscle function.

A functional class of genes known to impact ATP synthesis was also identified in our screen. This led us to discover that levamisole treatment decreased cellular ATP, which likely leads to metabolic exhaustion. Levamisole is commonly used by members of the *C. elegans* community to immobilize animals for imaging experiments. Thus, our results have broad significance as understanding the consequences of levamisole exposure is important for researchers who use this as a chemical anesthetizing agent.

## Acknowledgements

We thank and J. L. Bessereau for providing *unc-63(kr98-YFP)* and *krSi2(unc-49::tagRFP)*; other nematode strains used in this work were provided by the *Caenorhabditis* Genetics Center, which is funded by the NIH National Center for Research Resources (NCRR). We thank Todd Lamitina and Denis Touroutine for helpful discussions and critical reading of the manuscript. This work was supported by National Institute of Arthritis and Musculoskeletal Diseases (NIAMS) grant F32 AR060128 and National Institute of General Medical Science (NIGMS) IDeA Network of Biomedical Research Excellence (INBRE) P20 GM103446 Core Center Access Award (to J.E.T.).

## Footnotes

### Author Contributions

T.C. and J.T. conducted the screen. T.C., S.P., E.S., A.L., M.C., K.K. and J.T. performed time course assays. S.P. and E.M. carried out the confocal imaging. A.L. conducted the ATP quantitation. E.S., K.K. and J.T. carried out literature searches on identified genes. E.S. and J.T. created the figures. E.S., S.P. and J.T. wrote the manuscript. The authors declare no competing financial interests.

---

**Supplemental Table 1:**
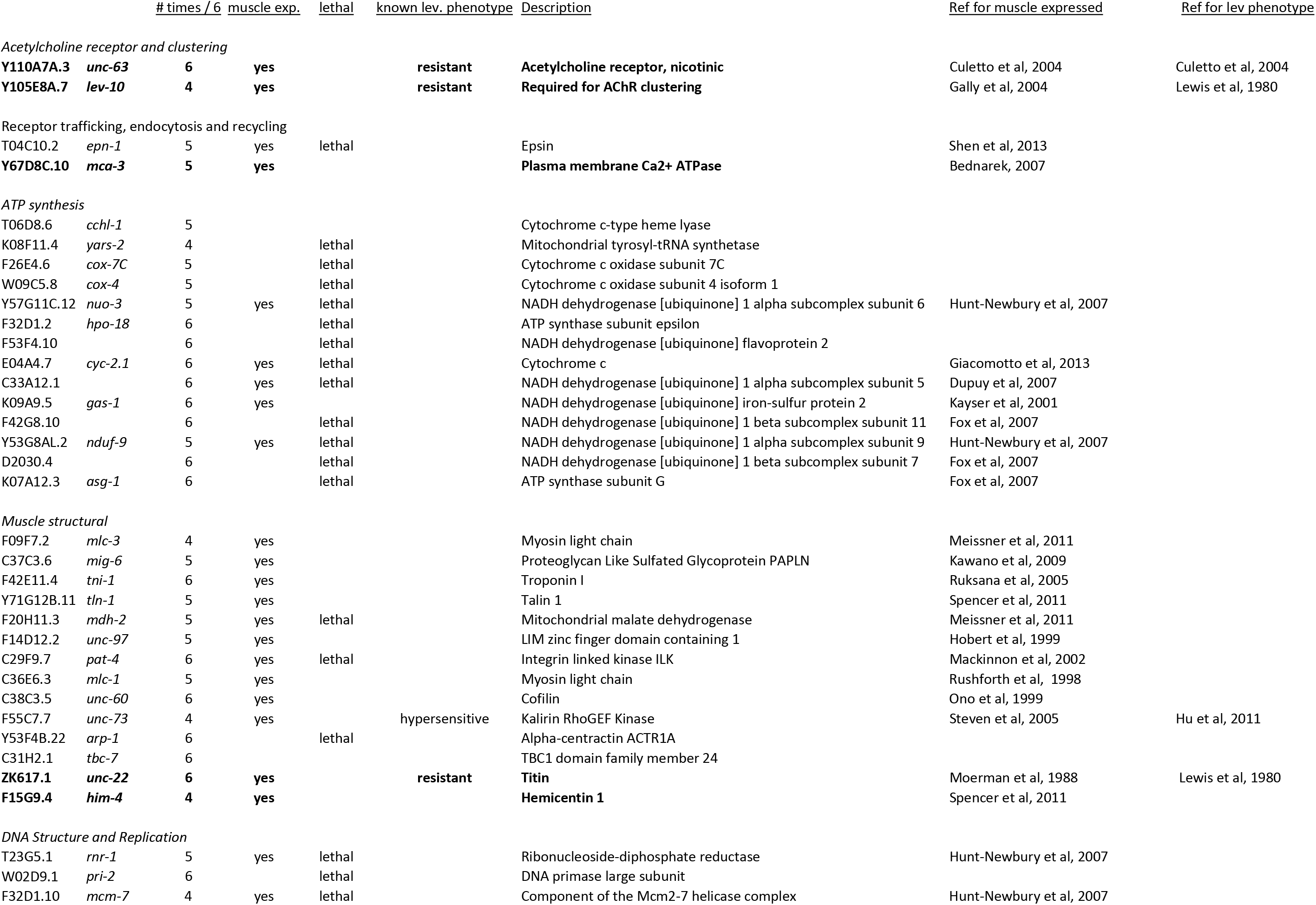
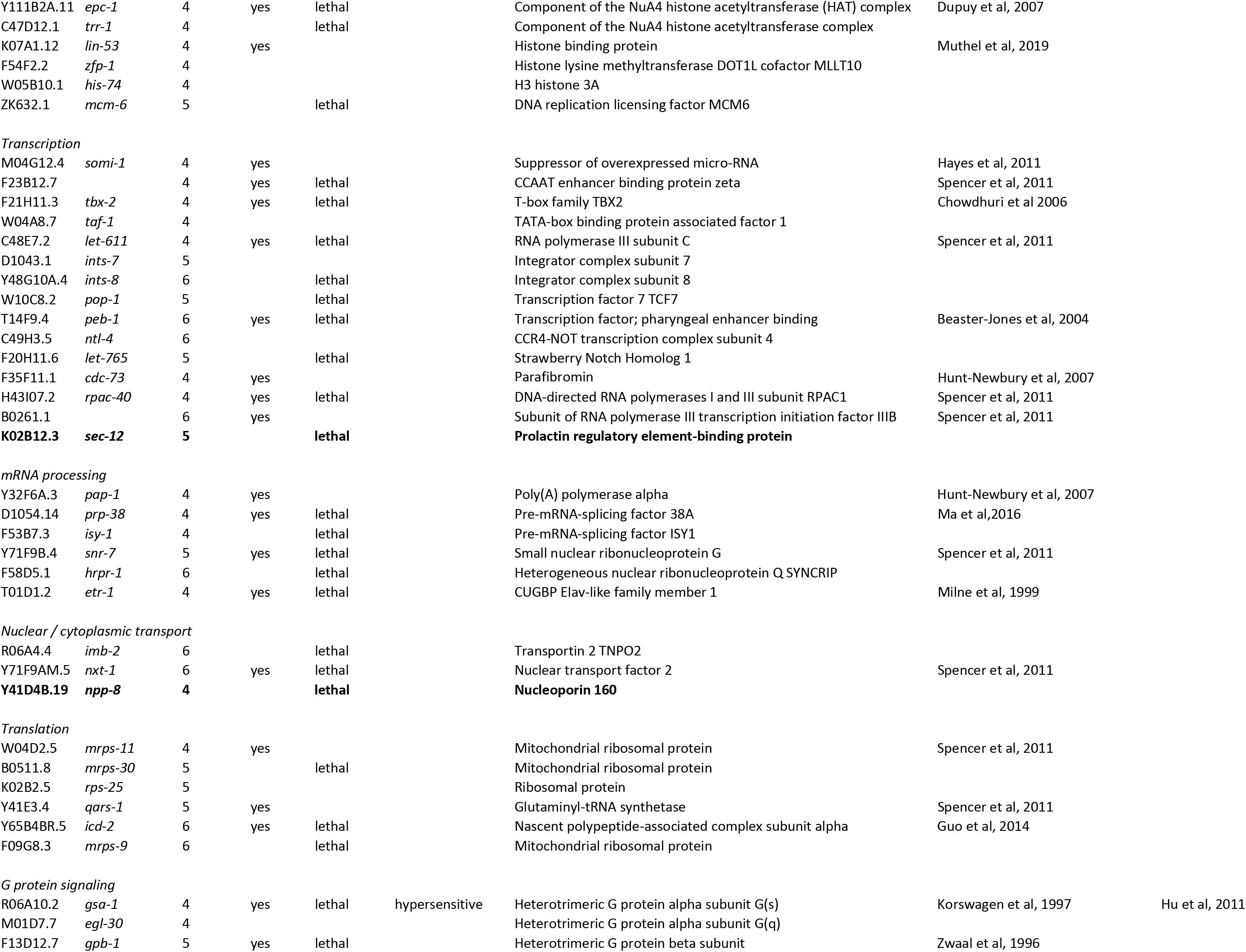
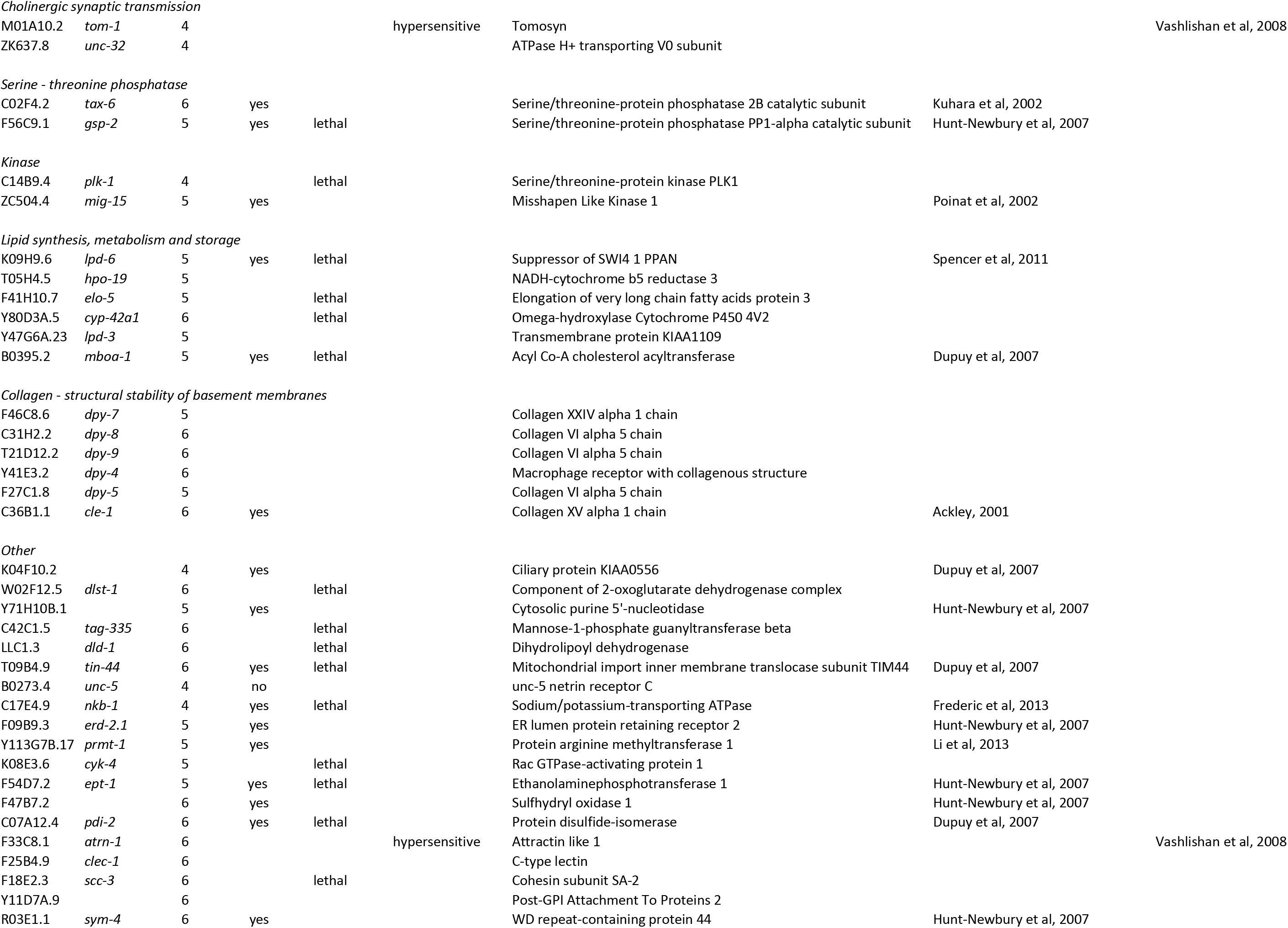
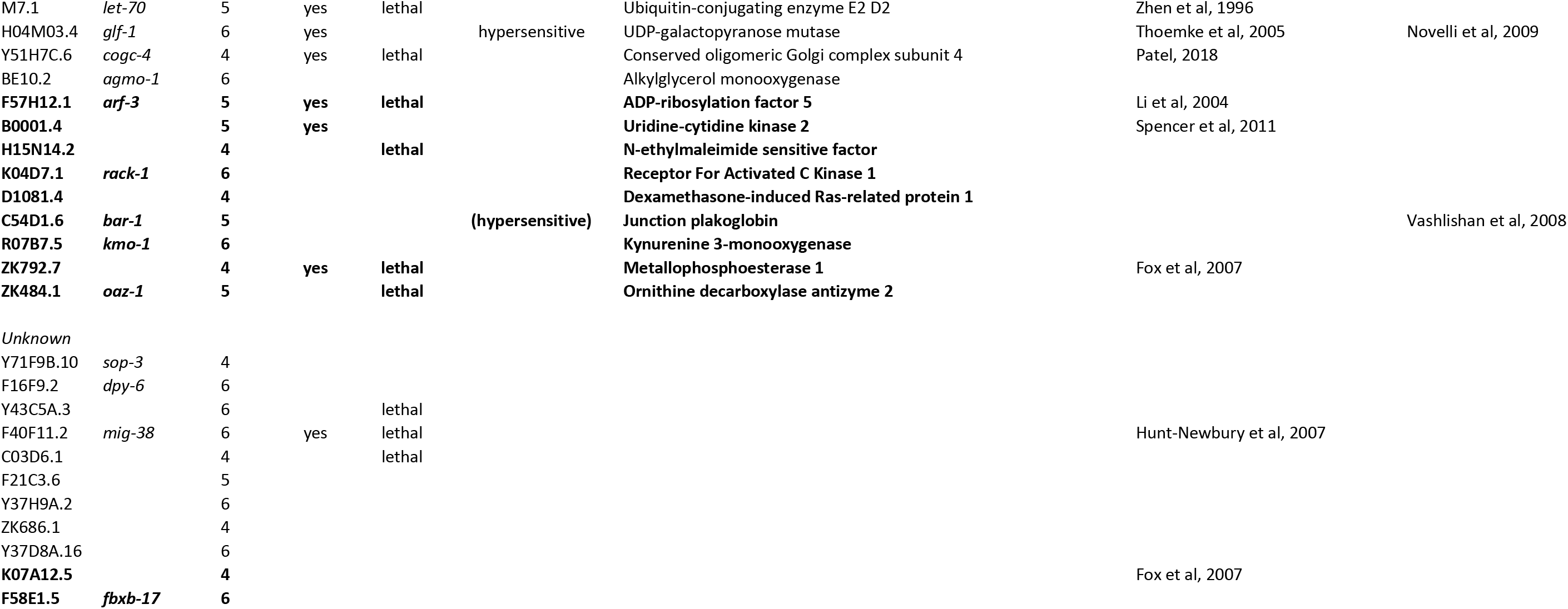
Genes identified in the RNAi screen for altered sensitivity to levamisole. Knockdown of the genes shown in **bold** caused levamisole resistance; all others resulted in hypersensitivity.

